# Temporal changes in the individual size distribution decouple long-term trends in abundance, biomass, and energy use of North American breeding bird communities

**DOI:** 10.1101/2022.11.08.515659

**Authors:** Renata M. Diaz, S. K. Morgan Ernest

## Abstract

**Aim:** A core objective of contemporary biodiversity science is to understand long-term trends in the structure and function of ecological communities. Different currencies of ecological function – specifically, total abundance, total standing biomass, and total metabolic flux – are naturally linked, but may become decoupled if the underlying size structure of a system changes. Here, we seek to establish how changes in community size composition modulate long-term relationships between different currencies of ecological function for North American birds.

**Location:** North America, north of Mexico.

**Time period:** 1988-2018.

**Major taxa studied:** Breeding birds.

**Methods:** We used species’ traits and allometric scaling to estimate individual size measurements and basal metabolic rate for birds observed in the North American Breeding Bird Survey. We compared the long-term trajectories for community-wide standing biomass and energy use to the long-term trends driven by changes in individual abundance alone. Finally, we used dissimilarity metrics to evaluate the link between changes in species and size composition and changes in the relationship between abundance- and size-driven dynamics.

**Results:** For a substantial minority of communities, shifts in community size composition have decoupled the long-term dynamics of biomass, energy use, and individual abundance. While trends in abundance were dominated by decreases, trends in biomass were evenly divided between decreases and increases, and trends in energy use featured more increases than expected given changes in abundance alone. Communities with decoupled dynamics showed greater increases in community-wide mean body size than other communities, but did not differ from other communities in overall turnover in species or size composition.

**Main conclusions:** Size- and abundance-based currencies of ecological function are linked, but not necessarily equivalent. For North American breeding birds, shifts in species composition favoring larger-bodied species may have partially offset declines in standing biomass driven by losses of individuals over the past 30 years.

## Introduction

Understanding the interrelated dynamics of size- and abundance-based dimensions of biodiversity is key to understanding biodiversity change in the Anthropocene. Total abundance - i.e. the total number of individual organisms present in a system - and size-based currencies - such as the total biomass or total metabolic flux (“energy use”) of a system - are intertwined, but not necessarily equivalent, measures of biological function. Abundance is more closely tied to species-level population dynamics, while size-based metrics more directly reflect assemblage-level resource use and contributions to materials fluxes at the broader ecosystem scale (Morlon et al. 2009, Dornelas et al. 2011, Connolly et al. 2005, White et al. 2007). While these currencies are naturally linked (Morlon et al. 2009, Henderson and Magurran 2010), changes in size composition can decouple the dynamics of one currency from another (Ernest et al. 2009, Dornelas et al. 2011, White et al. 2004, 2007, Yen et al. 2017). This can mean that intuition from one currency may be misleading about others; a trend in numerical abundance might mask alternative dynamics occurring with respect to biomass or total energy use (White et al. 2004). Moreover, changes in size composition strong enough to decouple currencies may be symptomatic of important changes in ecosystem status-e.g. abundance-biomass comparison curves (Petchey and Belgrano 2010) or size-biased extinctions (Young et al. 2016, Smith et al. 2018). This underscores the need to understand how these dynamics are playing out in the Anthropocene (Fisher et al. 2010).

Specifically, at the community scale, changes in the relationship between size and abundance can signal important shifts in community structure and functional composition. To the extent that size is a proxy for other functional traits, the dynamics of the community-level size structure (individual size distribution, ISD) over time may reflect processes related to niche structure (White et al. 2007, Petchey and Belgrano 2010). Strong size shifts can decouple the relationship between abundance and biomass. This is especially well-established in aquatic systems, where such changes in the scaling between abundance and biomass often signal ecosystem degradation (Warwick and Clarke 1994, Kerr and Dickie 2001, Petchey and Belgrano 2010). Compensatory shifts in the size structure can buffer community function (in terms of biomass or energy use) against changes in abundance (Ernest et al. 2009, White et al. 2004, Terry and Rowe 2015). Or, consistency in the size structure may maintain the relationship between size- and -abundance based currencies, even as species composition, total abundance, and total biomass and total energy use fluctuate over time, which can reflect consistency in the niche structure over time (Holling 1992).

It is important to improve our understanding of these dynamics for terrestrial animal communities in particular. In contrast to terrestrial trees and aquatic systems (Kerr and Dickie 2001, White et al. 2007), how the relationship between size and abundance changes over time, and the consequences of these changes for ecosystem-level properties, remain relatively unknown for terrestrial animals (but see White et al. (2004)). Terrestrial animal communities exhibit size structure (Thibault et al. 2011, Ernest 2005), and case studies have demonstrated that size shifts can decouple the dynamics of abundance, biomass, and energy use for terrestrial animals (White et al. 2004, Yen et al. 2017), but do not always do so (Hernández et al. 2011). Establishing generalities in these dynamics is especially pertinent in the Anthropocene, as these communities are experiencing extensive and potentially size-structured change, with implications at community, ecosystem, and global scales (Young et al. 2016, Schmitz et al. 2018).

Macroecological-scale synthesis on the interrelated dynamics of the ISD, total abundance, and community function for terrestrial animals has been constrained by 1) a lack of community-level size and abundance timeseries data for these systems (Thibault et al. 2011, White et al. 2007), and 2) appropriate statistical methods for relating change in the size structure to changes in abundance and function (Thibault et al. 2011, Yen et al. 2017). In contrast to aquatic and forest systems, most long-term surveys of animal communities do not collect data on individuals’ *sizes* across a full community (with the exception of small mammal studies, which have made major contributions to our understanding of the dynamics of size, abundance, and function for these systems; White et al. 2004, Ernest 2005, Hernández et al. 2011, Kelt et al. 2015). Global, continental, or population-wide studies capture different phenomena (White et al. 2007, McGill et al. 2015). The ISDs for terrestrial animals, and specifically for determinate growing taxa (e.g. mammals, birds), are often complex, multimodal distributions strongly determined by community *species* composition (Holling 1992, Thibault et al. 2011, Ernest 2005, Yen et al. 2017) and less statistically tractable than the power-law ISDs found in aquatic and tree systems (Kerr and Dickie 2001, White et al. 2007). As a result, we do not have a general understanding of either 1) how the size structures for these systems behave over time or 2) the extent to which changes in community size structure decouple the community-level dynamics of abundance, biomass, and energy use in these systems.

Here, we begin to address this gap by exploring how temporal changes in the size structure modulate the relationship between total abundance, energy, and biomass for communities of North American breeding birds. We used allometric scaling to estimate community size and abundance data for the North American Breeding Bird Survey, and evaluated how changes in total abundance, biomass, and energy use have co-varied from 1988-2018. Specifically, we examined: 1) How often do these currencies change together vs. exhibit decoupled dynamics?; 2) What are the dominant directions and magnitudes of the overall change over time and degree of decoupling between the currencies?; 3) To what extent do changes in species composition and community size structure translate into decoupling in the temporal trends of different currencies at the community scale?

## Methods

Code to replicate these analyses is available online on GitHub, and will be archived to Zenodo upon manuscript acceptance. For the purposes of double-blind review, an anonymized copy is available at https://github.com/bbssizeshifts/BBSsims.

### Bird abundance data

We used data from the Breeding Bird Survey (Pardieck et al. 2019) to evaluate trends in abundance, biomass, and energy use. The Breeding Bird Survey consists of roughly 40 kilometer-long survey routes distributed throughout the United States and Canada. Routes are surveyed annually during the breeding season (predominately May-June), via 50 3-minute point counts during which all birds seen or heard are identified to species (Pardieck et al. 2019).

Sampling began in 1966, and routes have been added over time to a current total of roughly 3000 routes (Pardieck et al. 2019) We explored trends in abundance, biomass, and energy use over the 30-year time period from 1989-2018. We selected these years to provide a temporal window sufficient to detect long-term trends (Cusser et al. 2020), while allowing for a substantial number of routes. To avoid irregularities caused by missing time steps, we restricted the main analysis to routes that had been sampled in at least 27 of 30 years in this window (n = 739) and compared these results to a more strict selection of routes that were sampled in every year (n = 199). Results for this more stringent subset of routes were qualitatively the same as for the more inclusive selection of routes (Appendix S1). We take the route to be the “community” scale (Thibault et al. 2011). We filtered the data to remove taxa that are poorly sampled through the point-count methods used in the Breeding Bird Survey, following Harris et al. (2018).

### Estimated size data

The Breeding Bird Survey dataset contains abundances for all species along each route in each year, but does not include measurements of individual body size. We generated body size estimates for individual birds assuming that intraspecific size distributions are normally distributed around a species’ mean body size (following Thibault et al. (2011)). Using records of species’ mean and standard deviation of body mass from Dunning (2008), we drew individuals’ body sizes from the appropriate normal distributions. For species for which there was not a standard deviation recorded in Dunning (2008) (185 species affected, of 421 total), we estimated the standard deviation using an allometric scaling relationship between mean and standard deviation in body mass constructed by fitting a linear model to the records that did have mean and standard deviation measurements. This resulted in the scaling relationship *log* (*variance*) – —5.273 + (*log*(*mass*) * 1.995)); model *R^2^* =.86; see also Thibault et al. (2011) for a similar scaling relationship. For species with multiple records in Dunning (2008), we averaged the mean and standard deviation across all records (averaging across sexes, subspecies, and records from different locations). We performed this averaging after estimating any missing standard deviation measurements. For each individual bird observed, we estimated metabolic rate as 10.5 * (*mass*^.713^) (Fristoe 2015, Nagy 2005, McNab 2009). For each route in a given year, we computed total energy use, total biomass, and total abundance by summing over all individuals observed on that route in that year. This method does not incorporate intraspecific variation in body size across geographies or over time (Dunning 2008, Gardner et al. 2011, Youngflesh et al. 2022). However, it makes it possible to conduct macroecological studies of avian size distributions at a spatial and temporal scale that would otherwise be impossible (Thibault et al. 2011).

### Comparing abundance- and size- based currencies

Comparing trends across different currencies is a nontrivial statistical problem. Because different currencies vary widely in their scale of measure (e.g. abundance in the hundreds of individuals; total biomass in the thousands of grams), it is challenging to interpret differences in magnitude of slope across different currencies. Transformation and scaling using common approaches (such as a square-root transformation, or rescaling each currency to a mean of 0 and a standard deviation of 1) destroys information about the degree of variability within each currency that is necessary in order to make comparisons *between* currencies for the same timeseries.

Therefore, rather than attempting to compare slopes across currencies or to transform different currencies to a common scale, we used a simple null model to generate the expected dynamics in biomass and energy use if the individual size distribution had remained constant over time, but allowed *abundance* to vary consistent with observed dynamics. In effect, we generated the expected dynamics of biomass and energy use if only abundance drove changes in those currencies over time. For each route, we characterized the “observed” timeseries of total biomass and total energy use by simulating size measurements for all individuals observed in each time step and summing across individuals, using the method described above. We then simulated timeseries for “abundance-driven” dynamics of biomass and energy use incorporating observed changes in community-wide abundance over time, but under a scenario of consistent species (and therefore approximate size) composition over time. For each community, we characterized the timeseries-wide probability of an individual drawn at random from the community belonging to a particular species (*P*(*s_i_*)) as each species’ mean relative abundance taken across all timesteps:

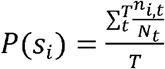

where *n_i,t_* is the abundance of species *i* in timestep *t, N_t_* is the total abundance of all species in timestep *t*, and *T* is the total number of timesteps. For each timestep *t*, we randomly assigned species’ identities to the total number of individuals of all species observed in that time step (*N_t_*) by drawing with replacement from a multinomial distribution with probabilities weighted according to *P*(*s*) for all species. We then simulated body size measurements for individuals, and calculated total energy use and total biomass, following the same procedure as for the observed community. This characterizes the dynamics for size-based currencies expected if the species (and size) composition of the community does not change over time, but incorporating observed fluctuations in total abundance.

### Long-term trends

For each route, we evaluated the “observed” 30-year trend in biomass (or energy use) and compared this to the trend derived from the “abundance-driven” null model using generalized linear models with a Gamma family and log link (appropriate for strictly-positive response variables such as biomass or total energy use). We fit four model formulas to characterize 1) the trend in biomass (or energy use) over time and 2) whether this trend deviates from the trend expected given only changes in individual abundance. These models correspond to qualitatively different “syndromes” of change:

1. *biomass ~year * dynamics* or *energy use ~year * dynamics* in which “dynamics” refers to being either the “observed” or “abundance-driven” (null model) dynamics. This model fits a slope and intercept for the observed trend in biomass or energy use over time, and a separate slope and intercept for the trend drawn from the abundance-driven, or null model, dynamics. We refer to this model as describing a syndrome of “Decoupled trends” between abundance-driven and observed dynamics.
2. *biomass ~year + dynamics* or *energy use ~year + dynamics*. This model fits a separate intercept, but not slope, for the abundance-driven and observed dynamics. This model was never selected as the best-performing description of community dynamics.
3. *biomass ~year* or *energy use ~year*. This model fits a temporal trend, but does not fit separate trends for the observed and abundance-driven dynamics. We refer to this syndrome as “Coupled trends” between abundance-driven and observed dynamics.
4. *biomass ~ 1* or *energy use ~ 1*. The intercept-only model describes no directional change over time for either the observed or abundance-driven dynamics, and we refer to this syndrome as describing “No directional change” for either type of dynamics.

We selected the best-fitting model using AICc. In instances where multiple models had AICc scores within two AICc units of the best-fitting model, we selected the simplest model within two AICc units of the best score.

For each route’s selected model, we extracted the predicted values for the first (usually 1988) and last (usually 2018) year sampled, for both the observed and null trajectories. We calculated the magnitude of change over time as the ratio of the last (2018) to the first (1988) value, and characterized the direction of the long-term trend as increasing if this ratio was greater than one, and decreasing if it was less than one.

### Relating change in community structure to decoupling between abundance and size-based dynamics

We used dissimilarity metrics to explore the extent to which change in community species or size composition caused decoupling between long-term trends in individual abundance and total biomass and energy use. These dissimlilarity metrics are most readily interpretable when making pairwise comparisons (as opposed to repeated comparisons over a timeseries). We therefore made comparisons between the first and last five-year intervals in each timeseries, resulting in a “begin” and “end” comparison separated by a relatively consistent window of time across routes (usually 19-20 years). The use of five-year periods corrects for sampling effects (White 2004), smooths out interannual variability, and, by including a relatively large proportion (1/3) of the total timeseries, partially mitigates the impact of scenarios where the start and end values do not align with the long-term trend.

We calculated three metrics to explore how changes in community composition and size structure translate into decoupling between abundance-driven and observed dynamics for biomass and energy use. First, we evaluated the change in average community-wide body size, calculated as the absolute log ratio of mean body size in the last five years relative to the mean body size in the first five years:

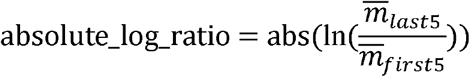

where 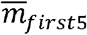 and 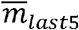 are the mean body size of all individuals observed in the first and last 5 years, respectively. Large changes in average body size are, by definition, expected to translate into decoupling between observed and abundance-driven dynamics.

Second, we calculated measures of turnover in the size structure and in species composition. We calculated turnover in the ISD using a measure inspired by an overlap measure that has previously been applied to species body size distributions in mammalian communities (Read et al. 2018). We characterized each “begin” or “end” ISD as a smooth probability density function by fitting a Gaussian mixture model (with up to 15 Gaussians, fit following Thibault et al. 2011) to the raw distribution of body masses, and extracting the fitted probability density at 1000 evaluation points corresponding to body masses encompassing and extending beyond the range of body masses present in this dataset (specifically, from 0 to 15 kilograms; mean body masses in this dataset range from 2.65 grams, for the Calliope hummingbird *Selasphorus calliope*, to 8.45 kg, for the California condor *Gymnogyps californianus*). We rescaled each density function such that the total probability density summed to 1. To calculate the degree of turnover between two ISDs, we calculated the area of overlap between the two density smooths as:

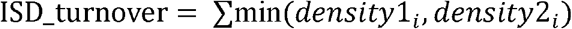

where densĩty1_*i*_ is the probability density from the density smooth for the first ISD at evaluation point *i*, and *density*2_*i*_ is the probability density from the density smooth for the second ISD at that evaluation point. We subtracted this quantity from 1 to obtain a measure of turnover between two ISDs.

To evaluate turnover in species composition between the five-year time periods, we calculated Bray-Curtis dissimilarity between the two communities using the R package *vegan* (Pinheiro et al. 2020).

We tested whether routes whose dynamics were best-described by each “syndrome” of change – i.e. “Decoupled trends”, “Coupled trends”, or “No directional change” – differed in 1) the magnitude of change in mean body size; 2) turnover in the ISD over time; or 3) species compositional turnover (Bray-Curtis dissimilarity) over time. For change in mean body size, we fit an ordinary linear model of the form *absolute_log_ratio ~ syndrome*. We used the absolute log ratio so as to focus on the magnitude, rather than the direction, of change in body size (see also Supp and Ernest (2014) for the use of the absolute log ratio to examine the magnitudes of differences between values). We compared this model to an intercept-only null model of the form *abs(log_ratio) ~ 1*. Because our metrics for turnover in the ISD and species composition are bounded from 0-1, we analyzed these metrics using binomial generalized linear models of the form *ISD_turnover ~ syndrome* and *Bray_Curtis_dissimilarity ~ syndrome*, and again compared these models to intercept-only null models. In instances where the model fit with a term for *syndrome* outperformed the intercept-only model, we calculated model estimates and contrasts using the R package *emmeans* (Lenth 2021).

## Results

Of the 739 routes in this analysis, approximately 70% (501/739 for biomass, and 509/739 for energy use) exhibited a significant temporal trend in either abundance or in biomass/energy use that resulted in the route being classified as exhibiting either “Decoupled trends” or “Coupled trends” (Table 1). All results were qualitatively the same using a subset of 199 routes with complete temporal sampling over time (Appendix S1). Trends driven by individual abundance, as reflected by the dynamics of a simple null model, were strongly dominated by declines (67% declines and 33% increases for abundance-driven dynamics in biomass, and 70% decreases and 30% increases for abundance-driven dynamics in energy use; Figure 2; Table 2). However, for biomass, the long-term temporal trends were evenly balanced between increases and decreases (49% increasing and 51% decreasing; Figure 2; Table 2). For energy use, there was a greater representation of decreasing trends than for biomass, but still less so than for strictly abundance-driven dynamics (65% decreasing and 35% increasing trends; Figure 2; Table 2).

**Figure 1.**
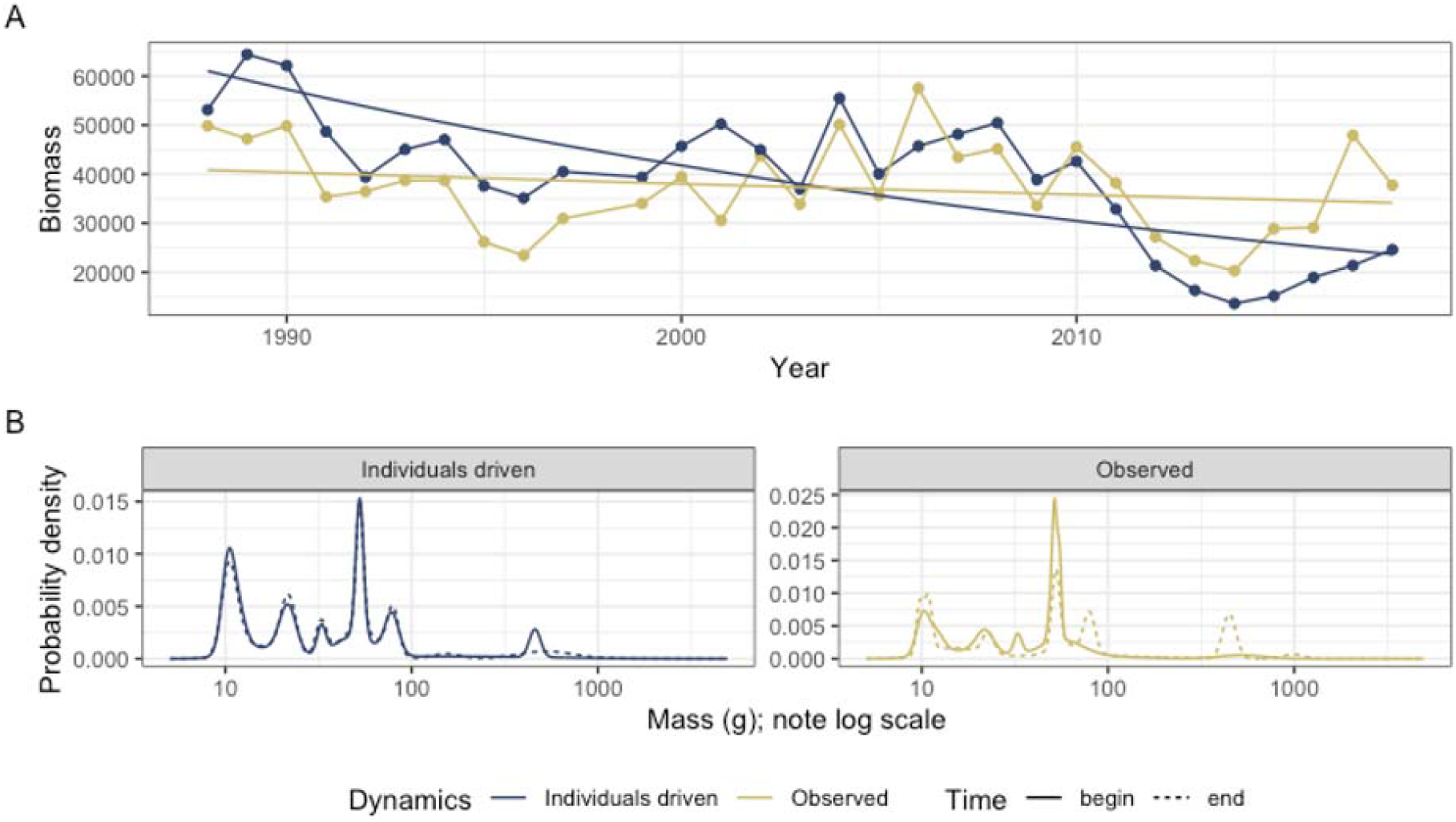
Illustration of abundance-driven (null model) dynamics as compared to observed dynamics (A), and the underlying dynamics of the ISD (B) for a sample route (Lindbrook, Alberta). **A. Dynamics of total biomass.** The gold points show the true values for total biomass in each year, and the blue points show the values for total biomass simulated from a null model that incorporates change in total abundance, but assumes no change in the size structure, over time. The smooth lines show the predicted values from a Gamma (log-link) linear model of the form *total_biomass ~ year * dynamics*, where *dynamics* refers to either the abundance-driven (null model) or observed dynamics. For this route, change in the individual size distribution has decoupled the dynamics of biomass from those that would occur due only to changes in abundance. The slope for abundance-driven dynamics is significantly more negative than for the observed dynamics (interaction term *p = 0.0013*) **B. Underlying changes in the ISD.** The individual size distributions for the first 5 years (solid lines) and last 5 years (dashed lines) of the timeseries. The x-axis is body size (as mass in grams; note log scale) and the y-axis is probability density from a Gaussian mixture model fit to a vector of simulated individual masses for all individuals observed in the years in questions, standardized to sum to 1. For the abundance-driven (blue) scenario, individuals’ species identities (which determine their body size estimates) are re-assigned at random weighted by each species’ mean relative abundance throughout the timeseries, resulting in a consistent individual size distribution over time. For the observed (gold) scenario, individuals’ body sizes are estimated based actual species abundances at each time step. For this route, species composition has shifted over time and produced different ISDs for the “begin” and “end” time periods. Specifically, the “end” ISD has peaks at larger body sizes (ca. 90g and 500g) not present in the “begin” ISD. This redistribution of density towards larger body sizes results in an overall increase in body size community wide, which partially offsets declines in total biomass from those expected given change in abundance alone.

**Figure 2.**
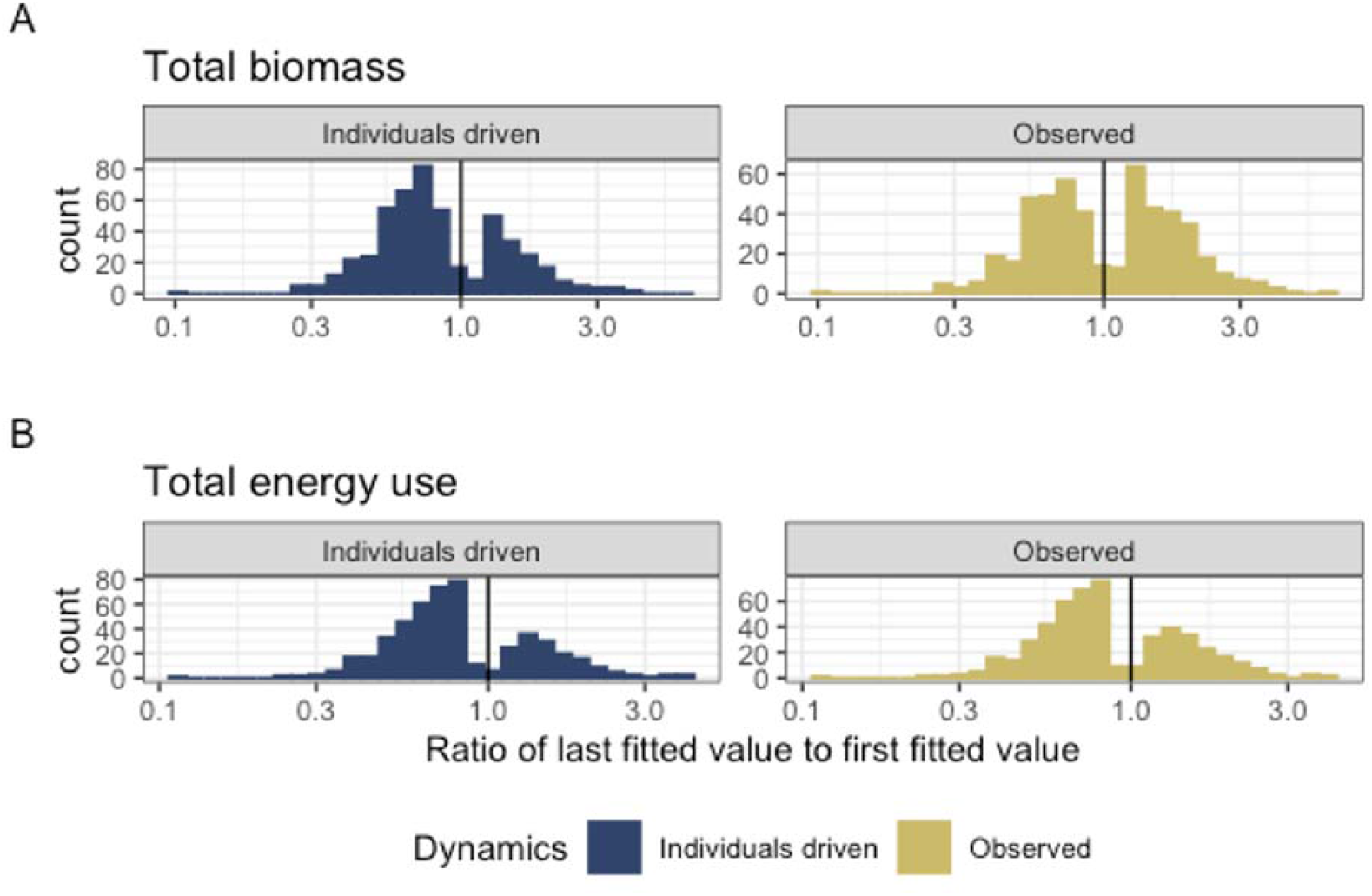
Histograms showing the direction and magnitude of long-term trends for the abundance-driven (null-model; left) and observed (right) changes in biomass (A) and energy use (B), for communities with a significant slope and/or interaction term (for biomass, 501/739 routes; for energy use, 509/739 routes; Table 1). Change is summarized as the ratio of the fitted value for the last year in the time series to the fitted value for the first year in the timeseries from the best-fitting model for that community. Values greater than 1 (vertical black line) indicate increases in total energy or biomass over time, and less than 1 indicate decreases. The abundance-driven dynamics (left) reflect the trends fit for the null model, while the observed dynamics (right) reflect trends incorporating both change in total abundance and change in the size structure over time. For communities best-described by syndromes of “coupled trends” or “no directional change”, the “abundance-driven” and “observed” ratios will be the same; for communities with “decoupled trends”, there will be different ratios for or “abundance-driven” and “observed” dynamics. Among routes with temporal trends (“coupled trends” or “decoupled trends”), there are qualitatively different continental-wide patterns in abundance-driven and observed dynamics for total biomass and total energy use. 70% of trends in abundance-driven (null model) dynamics for energy use are decreasing, and 67% for biomass (Table 2). For biomass, observed dynamics are balanced evenly between increases (49% of routes) and decreases (51%) - indicating that changes in the size structure produce qualitatively different long-term trends for biomass than would be expected given abundance changes alone. However, trends for energy use (which scales nonlinearly with biomass) are dominated by decreases (35% of routes), more closely mirroring the trends expected given changes in individual abundance alone.

**Table 1.** Table of the number and proportion of routes whose dynamics for total biomass and total energy use are best described by the following syndromes: no directional change (intercept- only model, *biomass ~ 1* or *energy use ~ 1*); a coupled trend (*biomass ~ year* or *energy use ~ year*); or a model with decoupled temporal trends for observed and abundance-driven dynamics (*biomass ~ year * dynamics* or *energy use ~ year * dynamics*, where *dynamics* refers to observed or null model, abundance-driven dynamics). 31-32% of routes are best described as syndromes of “No directional change” (intercept-only models). For the remaining routes, in most instances, the dynamics of biomass and energy use exhibit a temporal trend, but with no detectable difference in the temporal trends for abundance-driven and observed dynamics (“Coupled trends”). However, for a substantial minority of routes (20% overall for biomass, or 30% of routes with a temporal trend; 7% overall for energy use, or 10% of routes with a temporal trend), there is a detectable deviation between the trends expected due only to changes in abundance and the observed dynamics (“Decoupled trends”).

| Currency | Selected model | Number of routes | Proportion of routes |
| --- | --- | --- | --- |
| Total biomass | Intercept-only | 238 | 0.32 |
| Total biomass | Trend, not decoupled | 352 | 0.48 |
| Total biomass | Decoupled trend | 149 | 0.20 |
| Total energy use | Intercept-only | 230 | 0.31 |
| Total energy use | Trend, not decoupled | 456 | 0.62 |
| Total energy use | Decoupled trend | 53 | 0.07 |

**Table 2.** The proportion of trends that are increasing (specifically, for which the ratio of the last fitted value to the first fitted value > 1) for abundance-driven and observed dynamics, for routes exhibiting temporal trends (“coupled trends” or “decoupled trends”) in total biomass and total energy use (for biomass, *n* = 501; for energy use, *n =* 509). Trends that are not increasing are decreasing. Trends in abundance-driven dynamics are dominated by declines (67% of routes for total biomass, and 70% of routes for total energy). Observed dynamics for biomass differ qualitatively from the abundance-driven dynamics. Specifically, observed trends in biomass are evenly divided between increases and decreases (49% increasing). Observed trends in energy use more closely mirror abundance-driven trends (65% declines).

| Currency | Proportion of increasing abundance-driven trends | Proportion of increasing observed trends | Number of routes with temporal trends |
| --- | --- | --- | --- |
| Total biomass | 0.33 | 0.49 | 501 |
| Total energy use | 0.30 | 0.35 | 509 |

These divergent aggregate outcomes in individual abundance, energy use, and especially biomass occurred due to decoupling in the long-term trends for these different currencies. For a substantial minority of routes (20% of all routes for biomass, and 7% of all routes for energy use), long-term dynamics were best-described as a syndrome of “Decoupled trends” (that is, with a different slope for biomass or energy use-driven dynamics than for the “null”, individual abundance-driven, trend) (Table 1). When this decoupling occurred, it was dominated by instances in which the slope for individual abundance-driven dynamics was more negative than that for biomass or energy use (Figure 3).

**Figure 3.**
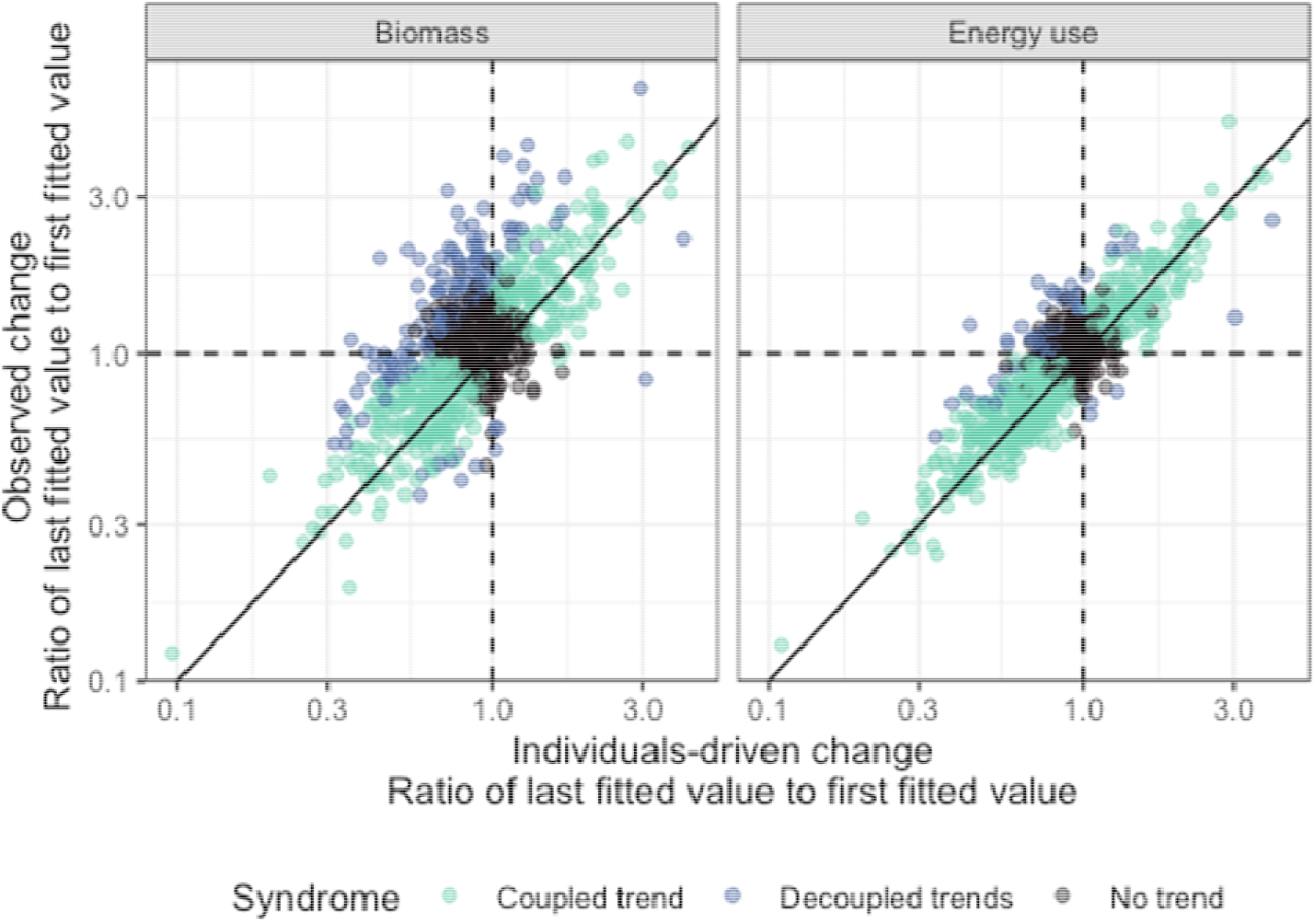
Observed change (ratio of last fitted value to first fitted value, y-axis) in total biomass (left) and total energy use (right) compared to the change expected only due to changes in individual abundance (ratio of last fitted value to first fitted value, x-axis), for all routes (*n* = 739) Values greater than 1 (dashed horizontal and vertical lines) mark positive (increasing) trends, while values less than 1 are negative trends. Each point marks the fitted values from a Gamma log-link generalized linear model of the form *biomass ~year * dynamics* for a given route, which calculates separate long-term slopes for observed and abundance-driven dynamics. Points are colored corresponding to the syndrome of change identified for each route. Deviations from the 1: 1 line (solid black line) reflect changes in the community size structure that modulate the relationship between total abundance and total biomass or energy use. Changes in total biomass and total energy use generally track changes driven by fluctuations in total abundance, with appreciable scatter around the 1:1 line. When this translates into a statistically detectable decoupling between observed and abundance-driven dynamics (a syndrome of “Decoupled trends”), this is usually in the form of abundance-driven change being more negative (a steeper decline or a smaller increase) than observed change in biomass or energy use (a less steep decline or larger increase), resulting in points falling above and to the left of the 1:1 line. This occurs more strongly and frequently for biomass than for energy use.

Decoupling between the long-term trajectories of individual abundance and energy use or biomass is, by definition, indicative of some degree of change in the ISD over time. Routes whose dynamics for biomass were best-described as syndromes of decoupled trends over time had a higher absolute log ratio of mean mass (i.e. greater magnitude of change, either increasing or decreasing, in mean mass over time) than routes with coupled or no directional trends (Figure 4; Appendix S2 Tables S1-S3). However, there was not a detectable difference in the degree of temporal turnover in the ISD overall (Figure 4; Appendix S2 Table S4), or in species composition (Figure 4; Appendix S2 Table S5), compared between routes that exhibited different syndromes of change.

**Figure 4.**
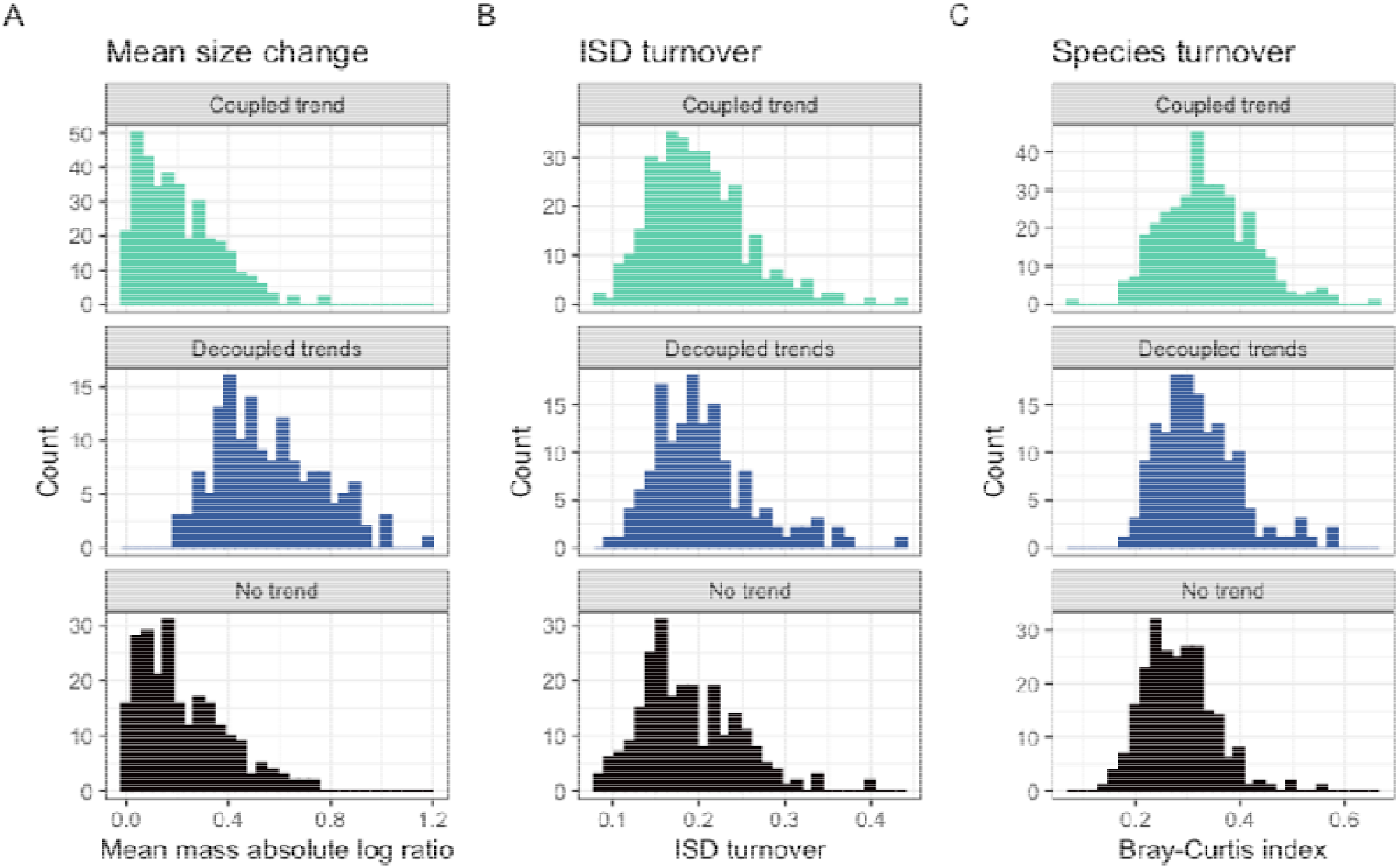
Histograms of (A) magnitude of change in mean body size from the first to the last five years of monitoring, (B) overall change in the size structure, and (C) change in species composition for routes whose dynamics for total biomass were best-described as syndromes of no directional change (bottom row; intercept-only model), decoupled trends for observed and abundance-driven dynamics (middle row), or coupled trends for observed and abundance-driven dynamics (top row). Change in mean body size (A) is calculated as the log of the ratio of the mean body size of all individuals observed in the last 5 years of the timeseries relative to the mean body size of all individuals observed in the first 5 years (see text). Overall change in the ISD (B) is calculated as the degree of turnover between the ISDs for the first and last five years of the timeseries (see text). Change in species composition (C) is characterized as Bray-Curtis dissimilarity comparing species composition in the first five years to the last five years. Routes that exhibit decoupling between observed and abundance-driven changes in total biomass exhibit high magnitudes of change in mean body size (middle row, panel A) compared to the changes seen in routes that show either no trend or “coupled” trends (see also Appendix 2 Tables S1-S3). However, routes with all three syndromes of dynamics (coupling, decoupling, or no trend) are not detectably different in the degree of overall change in the ISD or in species composition over time (panels B and C; Appendix 2 Tables S4 and S5).

## Discussion

### Abundance, biomass, and energy use are nonequivalent currencies

Simultaneously examining multiple currencies of community-level abundance revealed qualitatively different continent-wide patterns in the long-term trends for abundance in terms of individuals, biomass, and energy use. While long-term trends in individual abundance were dominated by decreases, long-term trends in biomass were evenly split between increases and decreases, and trends in energy use were again dominated by declines (Figure 2). These different currencies, though intrinsically linked, describe nonequivalent dimensions of community function and reflect different classes of structuring processes (Morlon et al. 2009). Abundance, in terms of individuals, is most directly linked to species-level population dynamics of the type often considered in classic, particularly theoretical, approaches to studying competition, compensation, and coexistence (e.g. Hubbell 2001; Chesson 2000). Biomass most directly reflects the productivity of a community and its potential contributions to materials fluxes in the broader ecosystem context, whereas energy use - by taking into account the metabolic inefficiencies of organisms of different body size - characterizes the total *resource use* of a community and may come the closest to capturing signals of bottom-up constraints, “Red Queen” effects, or zero-sum competitive dynamics (Van Valen 1973, Ernest et al. 2009, 2008, Morlon et al. 2009, White et al. 2004). Our results underscore that, while trends in abundance, biomass, and energy use naturally co-vary to some extent, shifts in the community size structure can and do produce qualitatively different trends for these different currencies. These may reflect contrasting long-term changes in different types of community processes - for example, shifts in habitat structure that affect the optimal body sizes for organisms in a system, but do not result in overall changes in resource availability (e.g. White et al. (2004)). Moreover, extrapolating the long-term trend from one currency to another may elide underlying changes in the community that complicate these dynamics. To appropriately monitor different dimensions of biodiversity change, it is therefore important to focus on the specific currency most closely aligned with the types of processes and dynamics - e.g. population fluctuations, resource limitation, or materials fluxes - of interest in a particular context.

### For North American breeding birds, biomass has declined less than abundance or energy use

For communities with a decoupling in the long-term trends of biomass, energy use, and abundance, this decoupling is indicative of a directional shift in the size structure of the community. For the communities of breeding birds across North America considered here, the long-term trends in total biomass are often less negative than trends in total abundance or total energy use (Figure 3), reflecting community-level increases in average body size that partially or completely buffer changes in total biomass against declines in abundance. This consistent (but not ubiquitous) signal contrasts with general, global concerns that larger-bodied organisms are more vulnerable to extinction and population declines than smaller ones (Young et al. 2016, Dirzo et al. 2014, Smith et al. 2018), but it is aligned with previous findings from the Breeding Bird Survey (Schipper et al. 2016). Because these data focus on *interspecific* variability in size, these findings reflect turnover in species composition broadly favoring larger-bodied species. A clear next step for this work is to identify the proximate and ultimate drivers of these shifts – for example, by identifying which groups or species are responsible for these trends, how these shifts vary over regions or habitat types, and if and how they are linked to underlying changes in habitat quality or anthropogenic disturbances. An equally important counterpoint to this work will be integrating potential shifts in *intraspecific* body size, and particularly declines in body size associated with rising temperatures, with the *interspecific* dynamics documented here (Youngflesh et al. 2022). In principle, declining body size over time could partially offset the interspecific size shifts observed here. Given that interspecific size shifts occur over a dramatically larger range of body sizes than do intraspecific size shifts, we anticipate that, in most instances, the community-wide dynamics would remain qualitatively despite intraspecific size change. While there are not currently robust estimates of intraspecific size changes at the scale of the full Breeding Bird Survey dataset, this is an excellent opportunity for a focused study in a well-documented system to elucidate the net effects of inter and intraspecific size change on community-level properties. Finally, we note that these increases in body size do not generally appear great enough to decouple the long-term trends in *energy use* from total abundance (Figure 3). Energy use scales nonlinearly with body size with an exponent less than 1, which means that community-wide increases in mean body size result in smaller increases in total energy use than in total biomass.

### Complex relationships between compositional change and community-level properties

The decoupling between the long-term trends for biomass, abundance, and energy use demonstrated in many of the communities studied here is symptomatic of a directional shift in the size structure - in these instances, generally favoring larger bodied species. However, examining the community-wide dynamics of turnover in species composition and the overall size structure reveals that the relationship between changes in community structure and changes in the scaling between different currencies of community-wide abundance is considerably more nuanced than simple directional shifts in mean size. Routes that exhibit a statistically detectable decoupling between total biomass and total abundance show large changes in average body size compared to routes for which biomass and abundance either change more nearly in concert with each other or do not show temporal trends (Figure 4; Appendix 2 Tables S1-S3). This aligns naturally with mathematical intuition given the intrinsic relationship between average body size, total abundance, and total biomass. However, these routes are *not* extraordinary in terms of their overall degree of temporal turnover in either the size structure or in species composition. Rather, the levels of turnover in overall community structure are comparable between routes that show decoupling between abundance and biomass, statistically indistinguishable trends, or no temporal trends in either currency (Figure 4; Appendix 2 Tables S4-S5).

For many communities, therefore, there has been appreciable change in the species and size composition that simply does not manifest in a shift in the overall community-wide mean body size or mean metabolic rate sufficient to decouple the dynamics of biomass, abundance, and energy use. These changes may signal changes in functional composition equally important as the ones that manifest in directional shifts in community-wide average body size. For the complex, multimodal size distributions that are the norm for avian communities (Thibault et al. 2011), changes in the number and position of modes may be as important as changes in higher- level statistical moments such as the overall mean. At present, the field lacks the statistical tools and conceptual frameworks to quantify and interpret these nuanced changes, especially at the macroecological scale of the current study (Thibault et al. 2011, Yen et al. 2017). However, this is an excellent opportunity for more system-specific work, informed by natural history knowledge and process-driven expectation, to characterize more nuanced changes in the size structure of specific communities and identify the underlying drivers of these changes. To facilitate these efforts in the context of the Breeding Bird Survey, we are developing a R package to characterize the individual size distributions for avian communities based on species’ identities and/or mean body sizes.

### Conclusion

This analysis demonstrates the current power, and limitations, of a data-driven macroecological perspective on the interrelated dynamics of community size structure and different dimensions of community-wide abundance for terrestrial animal communities. For breeding bird communities across North America, we find that changes in species and size composition produce qualitatively different aggregate patterns in the long-term trends of abundance, biomass, and energy use, highlighting the nuanced relationship between these related, but decidedly nonequivalent, currencies and reflecting widespread changes in community size structure that may signal substantive changes in functional composition. Simultaneously, the complex relationship between turnover in community species and size composition, and the scaling between different currencies of community-level abundance, highlights opportunities for synergies between recent computational and statistical advances, case studies grounded in empiricism and natural history, and future macroecological-scale synthesis to realize the full potential of this conceptual space.

## Data availability

All data and code supporting this manuscript are available online on GitHub at https://www.github.com/diazrenata/BBSsims. For the purposes of double-blind review, we have uploaded an identical copy of these materials to a temporary GitHub repository at https://github.com/bbssizeshifts/BBSsims. Upon manuscript acceptance, they will be archived in perpetuity on Zenodo.

## Ethics approval, patient consent, reproduction of material, and clinical trial registration

These do not apply to this manuscript.

## Biosketches

RMD is a computational biodiversity scientist studying how the emergent properties of ecological systems respond to global change. SKME is an ecologist who studies long-term changes in communities and the processes driving those changes.

## Data availability

All data and code supporting this manuscript are available online on GitHub. For the purposes of double-blind review, we have uploaded a copy of these analyses to a temporary GitHub repository at https://github.com/bbssizeshifts/BBSsims. Upon manuscript acceptance, these will be archived in perpetuity on Zenodo.

