## Appendix S1 for "Temporal changes in the individual size distribution decouple long-term trends in abundance, biomass, and energy use of North American breeding bird communities"

Figures and tables from the main analysis, restricted to 199 routes with perfect temporal coverage (i.e. no missing time steps). All results are qualitatively the same as for the main analysis (739 routes, with a minimum of 27 of 30 time steps sampled for each route).

Table of Contents

### Appendix S1 Figure S1.

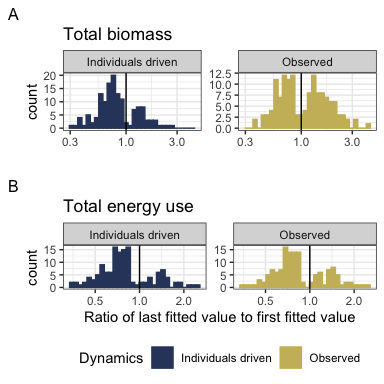

**Appendix S1 Figure S1**. Histograms showing the direction and magnitude of long-term trends for the individuals-driven (null-model; left) and observed (right) changes in biomass (A) and energy use (B), for communities with a significant slope and/or interaction term (for biomass, 141/199 routes; for energy use, 137/199 routes; Table 1). Change is summarized as the ratio of the fitted value for the last year in the time series to the fitted value for the first year in the timeseries from the best-fitting model for that community. Values greater than 1 (vertical black line) indicate increases in total energy or biomass over time, and less than 1 indicate decreases. The individuals-driven dynamics (left) reflect the trends fit for the null model, while the observed dynamics (right) reflect trends incorporating both change in total abundance and change in the size structure over time. For communities best-described by syndromes of “coupled trends” or “no directional change”, the “individuals-driven” and “observed” ratios will be the same; for communities with “decoupled trends”, there will be different ratios for or “individuals-driven” and “observed” dynamics.

Among routes with temporal trends (“coupled trends” or “decoupled trends”), there are qualitatively different continental-wide patterns in individuals-driven and observed dynamics for total biomass and total energy use. 76% of trends in individuals-driven (null model) dynamics for energy use are decreasing, and 72% for biomass (Table 2). For biomass, observed dynamics are balanced evenly between increases (50% of routes) and decreases (50%) - indicating that changes in the size structure produce qualitatively different long-term trends for biomass than would be expected given abundance changes alone. However, trends for energy use (which scales nonlinearly with biomass) are dominated by decreases (69% of routes), more closely mirroring the trends expected given changes in individual abundance alone.

### Appendix S1 Figure S2

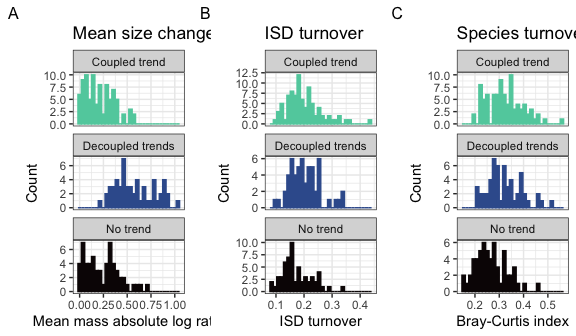

**Appendix S1 Figure S2.** Histograms of (A) change in mean body size from the first to the last five years of monitoring, (B) overall change in the size structure, and (C) change in species composition for routes whose dynamics for total biomass were best-described using no temporal trend (bottom row; intercept-only model), separate trends for observed and individuals-driven dynamics (middle row), or the same trend for observed and individuals-driven dynamics (top row). Change in mean body size (A) is calculated as the ratio of the mean body size of all individuals observed in the last 5 years of the timeseries relative to the mean body size of all individuals observed in the first 5 years. Overall change in the ISD (B) is calculated as the degree of turnover between the ISDs for the first and last five years of the timeseries (see text). Change in species composition (C) is Bray-Curtis dissimilarity comparing species composition in the first five years to the last five years.

### Appendix S1 Table S1.

| Currency | Selected model | Number of routes | Proportion of routes |
| --- | --- | --- | --- |
| Total biomass | Intercept-only | 58 | 0.29 |
| Total biomass | Trend, not decoupled | 86 | 0.43 |
| Total biomass | Decoupled trend | 55 | 0.28 |
| Total energy use | Intercept-only | 62 | 0.31 |
| Total energy use | Trend, not decoupled | 115 | 0.58 |
| Total energy use | Decoupled trend | 22 | 0.11 |

**Appendix S1 Table S1**. Table of the number and proportion of routes whose dynamics for total biomass and total energy use are best described by the following syndromes: no directional change (intercept-only model, biomass ~ 1 or energy use ~ 1); a coupled trend (biomass ~ year or energy use ~ year); or a model with decoupled temporal trends for observed and individuals-driven dynamics (biomass ~ year * dynamics or energy use ~ year * dynamics, where dynamics refers to observed or null model, individuals-driven dynamics).

### Appendix S1 Table S2

| Currency | Proportion of increasing individuals-driven trends | Proportion of increasing observed trends | Number of routes with temporal trends |
| --- | --- | --- | --- |
| Total biomass | 0.28 | 0.50 | 141 |
| Total energy use | 0.24 | 0.31 | 137 |

*Appendix S1 Table S2*. The proportion of trends that are increasing (specifically, for which the ratio of the last fitted value to the first fitted value > 1) for individuals-driven and observed dynamics, for routes exhibiting temporal trends (“coupled trends” or “decoupled trends”) in total biomass and total energy use. Trends that are not increasing are decreasing.

### Appendix S1 Table S3.

| Res.Df | RSS | Df | Sum of Sq | F | Pr(>F) |
| --- | --- | --- | --- | --- | --- |
| 196 | 5.993024 | NA | NA | NA | NA |
| 198 | 11.105203 | -2 | -5.11218 | 83.59613 | 0 |

**Appendix S1 Table S3**. ANOVA table comparing ordinary linear models of the form abs_log_ratio ~ syndrome and abs_log_ratio ~ 1.

### Appendix S1 Table S4.

| categorical_fit | emmean | SE | df | lower.CL | upper.CL |
| --- | --- | --- | --- | --- | --- |
| Coupled trend | 0.2084997 | 0.0188558 | 196 | 0.1713133 | 0.2456861 |
| Decoupled trends | 0.5779023 | 0.0235784 | 196 | 0.5314024 | 0.6244021 |
| No trend | 0.2385438 | 0.0229605 | 196 | 0.1932625 | 0.2838251 |

**Appendix S1 Table S4.** Estimates (calculated using emmeans (Lenth 2021)) for the mean absolute log ratio of mean mass for routes whose dynamics for biomass were best-described by different syndromes of change.

### Appendix S1 Table S5

| contrast | estimate | SE | df | t.ratio | p.value |
| --- | --- | --- | --- | --- | --- |
| Coupled trend - Decoupled trends | -0.3694026 | 0.0301908 | 196 | -12.235620 | 0.0000000 |
| Coupled trend - No trend | -0.0300441 | 0.0297107 | 196 | -1.011221 | 0.5706639 |
| Decoupled trends - No trend | 0.3393585 | 0.0329108 | 196 | 10.311453 | 0.0000000 |

**Appendix S1 Table S5**. Contrasts for absolute log ratio of mean mass, calculated using emmeans (Lenth 2021).

### Appendix S1 Table S6

| Resid. Df | Resid. Dev | Df | Deviance | Pr(>Chi) |
| --- | --- | --- | --- | --- |
| 196 | 4.053082 | NA | NA | NA |
| 198 | 4.253015 | -2 | -0.1999328 | 0.9048678 |

*Appendix S1 Table S6*. ANOVA table comparing binomial generalized linear models of the form ISD_turnover ~ syndrome and ISD_turnover ~ 1.

### Appendix S1 Table S7.

| Resid. Df | Resid. Dev | Df | Deviance | Pr(>Chi) |
| --- | --- | --- | --- | --- |
| 196 | 4.455349 | NA | NA | NA |
| 198 | 5.044039 | -2 | -0.5886892 | 0.7450197 |

**Appendix S1 Table S7**. ANOVA table comparing binomial generalized linear models of the form Bray_Curtis_dissimilarity ~ syndrome and Bray_Curtis_dissimilarity ~ 1.
