## Appendix S2 for "Temporal changes in the individual size distribution decouple long-term trends in abundance, biomass, and energy use of North American breeding bird communities"

Statistical comparisons of distributions in Figure 4.

Table of Contents

### Appendix S2 Table S1.

| Res.Df | RSS | Df | Sum of Sq | F | Pr(>F) |
| --- | --- | --- | --- | --- | --- |
| 736 | 20.81904 | NA | NA | NA | NA |
| 738 | 35.42466 | -2 | -14.60562 | 258.1708 | 0 |

*Appendix S2 Table S1.* ANOVA table comparing ordinary linear models of the form abs_log_ratio ~ syndrome and abs_log_ratio ~ 1. The fit incorporating syndrome is superior to the intercept-only model (p < 0.0001).

### Appendix S2 Table S2

| categorical_fit | emmean | SE | df | lower.CL | upper.CL |
| --- | --- | --- | --- | --- | --- |
| Coupled trend | 0.2012914 | 0.0089644 | 736 | 0.1836926 | 0.2188902 |
| Decoupled trends | 0.5587675 | 0.0137784 | 736 | 0.5317179 | 0.5858171 |
| No trend | 0.2203741 | 0.0109019 | 736 | 0.1989715 | 0.2417766 |

**Appendix S2 Table S2.** Estimates (calculated using emmeans (Lenth 2021)) for the mean absolute log ratio of mean mass for routes whose dynamics for biomass were best-described by different syndromes of change. Routes with decoupled long-term trends between biomass and individuals-driven dynamics have higher absolute log ratios (mean .56, 95% credible interval .53-.58) than routes with covarying trends in biomass and individual abundance (mean of .2; 95% interval .18-.22) or no detectable temporal trend (mean of .22; .2-.24).

### Appendix S2 Table S3

| contrast | estimate | SE | df | t.ratio | p.value |
| --- | --- | --- | --- | --- | --- |
| Coupled trend - Decoupled trends | -0.3574762 | 0.0164379 | 736 | -21.747096 | 0.0000000 |
| Coupled trend - No trend | -0.0190827 | 0.0141142 | 736 | -1.352018 | 0.3669124 |
| Decoupled trends - No trend | 0.3383935 | 0.0175697 | 736 | 19.260017 | 0.0000000 |

**Appendix S2 Table S3**. Contrasts for absolute log ratio of mean mass, calculated using emmeans (Lenth 2021). There is a significant contrast between routes with decoupled trends and the other two syndromes of dynamics (both contrasts, p < 0.001), but not between routes showing the “no trend” and “coupled trend” syndromes (contrast p = .37).

### Appendix S2 Table S4

| Resid. Df | Resid. Dev | Df | Deviance | Pr(>Chi) |
| --- | --- | --- | --- | --- |
| 736 | 14.09240 | NA | NA | NA |
| 738 | 14.28236 | -2 | -0.1899672 | 0.9093878 |

**Appendix S2 Table S4**. ANOVA table comparing binomial generalized linear models of the form ISD_turnover ~ syndrome and ISD_turnover ~ 1. The model incorporating syndrome is not superior to the intercept-only model (p = .91).

### Appendix S2 Table S5

| Resid. Df | Resid. Dev | Df | Deviance | Pr(>Chi) |
| --- | --- | --- | --- | --- |
| 736 | 20.09178 | NA | NA | NA |
| 738 | 22.11983 | -2 | -2.028057 | 0.3627546 |

**Appendix S2 Table S5**. ANOVA table comparing binomial generalized linear models of the form Bray_Curtis_dissimilarity ~ syndrome and Bray_Curtis_dissimilarity ~ 1. The model incorporating syndrome is not superior to the intercept-only model (p = .36).
